# No atypical white-matter structures in grapheme- or color-sensitive areas in synesthetes

**DOI:** 10.1101/618611

**Authors:** Franziska Weiss, Mark W. Greenlee, Gregor Volberg

**Affiliations:** University of Regensburg, Germany

**Keywords:** synesthesia, grapheme, cross-activation theory, DTI

## Abstract

Grapheme-color synesthetes experience colors when presented with written language characters. In this study diffusion-weighted imaging was used to investigate white matter alterations in color-and grapheme-processing brain areas in synesthetes as a possible factor for the color sensations. Regions of interest were defined by means of neuroanatomical atlantes, functional localizer tasks and retinotopic mapping. None of the regions showed differences in white matter structure between synesthetes and a control population, as revealed by fractional anisotropy and mean diffusivity measures. Quite the contrary, the data broadly supported the null hypothesis of no group differences in white matter microstructure. This finding is in line with recent studies suggesting no atypical neuroanatomy in grapheme-color synesthetes.

## 1. Introduction

The phenomenon known as Synesthesia has been known to both medical professionals as well as scientists for decades. In 1913, Coriat described the case of a patient who associated certain words, names or sounds with various shades of the color blue. These sensations persisted since childhood, were unidirectional in nature and remained stable over time. Coriat drew the conclusion that the phenomenon must be a “cortical one, possibly physiological”, and that synesthesia should arise from “a congenital defect of the nervous system, in which the stimulation of one center passes over into another” (Coriat, 1913). Today, this would be known as auditory-color synesthesia, and his theory on the origins of the synesthetic experience as a version of the cross-activation theory (Ramachandran & Hubbard, 2001).

Although an increasing amount of research has been done to explain the origins of the synesthetic experience, no clear conclusion has been drawn as to why it occurs. A model specifically addressing grapheme-color synesthesia is the cross-activation theory by Ramachandran and Hubbard (2001; Hubbard, Brang, & Ramachandran, 2011). This theory suggests that grapheme-color synesthetes exhibit increased neural connections between color processing areas and grapheme processing areas. Due to this cross-wiring, the neural representation of a grapheme automatically excites connected color processing neurons and, consequently, leads to a percept of a colored grapheme. For the brain areas involved in this cross-activation, the visual word form area (VWFA) and the visual area 4 (V4) have been proposed. Functional and neuropsychological data show that these structures are specialized for grapheme and color processing, respectively (Roe et al., 2012; Bartolomeo, Bachoud-Lévi, & Thiebaut de Schotten, 2014; McCandliss, Cohen, & Dehaene, 2003). They are located in juxtaposition in the inferotemporal cortex.

The main prediction of the cross-activation theory is an atypical brain anatomy in synesthetes, specifically in those brain areas where the synesthesia-inducing stimulus (‘inducer’) and the concurrent synesthetic sensation (‘concurrent’) are processed. Hence, grapheme-color synesthetes should hold a structural hyperconnectivity between VWFA and V4. Structural brain anatomy in synesthetes has been studied measuring gray matter volume with voxel-based morphometry (VBM). This method allows for investigation of cortical density. Increased gray matter volume in synesthetes’ color-processing areas was found in some studies using a-priori defined ROIs, but not on a whole-brain analysis level (Banissy et al., 2012; Hupé & Dojat, 2015; Jäncke, Beeli, Eulig, & Hänggi, 2009; McErlean, Janik McErlean, & Banissy, 2017; Weiss & Fink, 2009). Rouw and Scholte (2010) found that gray-matter volume differences co-varied with the subjective quality of the synesthetic perception across synesthetes. Some synesthetes experience synesthetic color as occurring in the outside world (‘projectors’), while others experience the colors as internal sensations (‘associators’). The former showed increased gray matter volume in V1, while the latter exhibited increased gray matter volume in hippocampus. Both types of synesthetes had increased gray matter volume in the superior parietal cortex (Rouw & Scholte, 2010). A recent study revealed increased gray matter in the right amygdala, left cerebellum and the left angular gyrus in synesthetes who report additional experiences accompanying number perception (Arend, Yuen, Sagi, & Henik, 2018). While these findings suggest neural alterations in synesthetes, the findings are inconclusive with respect to the affected structures and, consequently, are not decisive for the cross-activation theory.

Another method for investigating neurostructural differences is diffusion-weighted imaging (DWI), which calculates the diffusion of water molecules in different tissues (Van Hecke, Emsell, & Sunaert, 2015). The diffusion tensor model (DTI) provides diffusion parameters like fractional anisotropy (FA) and mean diffusivity (MD). FA is a parameter for the directionality of diffusion within a voxel and, hence, can be regarded as a measure of coherent fibre tracts in the white matter, dense axonal packing and high myelination. Increased FA values in synesthetes compared to controls could therefore be interpreted as more white matter and more connections in the regarding area. MD, on the other hand, shows the overall diffusivity or water content within an voxel (Van Hecke et al., 2015). The more restricted tissue a voxel contains, e.g. neuronal fibers with myelination, the smaller MD will be. Thus, it can be regarded as an inverse measure of tissue density.

The first direct indication of enhanced structural connectivity in synesthetes as measured by FA was found by Rouw and Scholte (2007). They found increased FA values for synesthetes in the right inferior temporal cortex, the left superior parietal and the superior frontal cortex. This finding could not be replicated to date (Hupé, Bordier, & Dojat, 2012; Jäncke et al., 2009). One source for methodological problems might be Tract-Based Spatial Statistics (TBSS; Smith et al., 2006) as applied in the original study. In this procedure, a skeleton is created according to thinned main tracts of the mean FA image across subjects. The generated skeletons don’t underlie the random field theory, which might have led to false positives in their analysis approach (see also Hupé & Dojat, 2015). Further, the non-linear registration of individual diffusion images to a standard space and reduction of the FA values to a skeleton of main tracts goes along with loss of individual diffusion information. While this method might be interesting for an exploratory analysis, it is not sufficient for the detection of differences in specific areas, as there are *a priori* defined ROIs proposed by the cross-activation theory.

Based on a critical overview and new data, Dojat et al. (2018) recently made the point that there are no structural differences between the synesthetic and the non-synesthetic brain. In a sample of 22 synesthetes and 25 controls measured with DTI along sixty isotropically different directions, only small to no group differences in FA could be found. Unlike in the previously mentioned studies, the authors also examined MD as an inverse measure for tissue density. This parameter did not significantly differ between groups either. However, as in the previously mentioned studies, FA differences were measured along a skeleton using TBSS.

The mapping of FA and MD to a standard brain or mean skeleton as seen in Dojat, Pizzagalli and Hupe (2018) and Rouw and Scholte (2007) leads to a loss of individual diffusion information. In this study, we present a different approach to the question of microstructural differences in color and letter processing areas in the brain of synesthetes. In contrast to the above mentioned studies, we examined the diffusion parameters FA and MD within the individual head space. Furthermore, we did not search for group differences on an exploratory whole brain level, which needs a larger sample size to reach full statistical power, but concentrated the investigation on the areas deemed critical for synesthetic color sensations by proponents of the cross-activation theory. To account for potentially misplaced ROIs, multiple methods were used to define color and letter processing areas anatomically and functionally. This also allows to compare our data to different studies using different approaches of ROI definition, as well as comparison between anatomical and functional forming of V4 and VWFA.

As a further advancement, Bayesian inference was used for hypothesis testing in this study besides Welch’s t-test. One practical advantage of Bayesian testing is that it allows the accumulation of evidence for a null hypothesis and for the alternative hypothesis at the same time. Moreover, because the BF is a relative quantity, it does not overstate the evidence for H_0_with low sample sizes (Morey & Rouder, 2011a). The Bayesian framework is therefore well suited for assessing null effects.

The inference reveals Bayesian factors (BFs) for any given test that indicate the relative posterior likelihood for two complementary hypotheses, H_0_and H_1_, that is obtained during data collection. For example, BF_10_= 6 means that the likelihood for H_1_is 6 times higher than for H_0_given the acquired data. BFs of 1 −3 mark anecdotal evidence, 3 −10 moderate evidence, and 10 −30 strong evidence in favor of the H_1_(e. g. Lee & Wagenmakers, 2014). The reciprocal values mark corresponding evidence in favor of the H_0_.

## 2. Methods

### 2.1 Participants

We obtained functional and structural MRI data from 11 synesthetes and 12 matched non-synesthetic control participants. The mean age was 22.5 years (19-30) in the synesthetic group and 25.5 years (20-39) in the control group. In each group, two of the participants were male. One of the synesthetes was left handed, the rest of the participants were right handed.

Synesthetes were recruited via written and oral announcements at the University of Regensburg and via online announcements in synesthesia social media groups. Interested synesthetes completed an online test battery (Eagleman, Kagan, Nelson, Sagaram, & Sarma, 2007) prior to the study. The test battery includes a synesthetic color picker task, in which participants have to choose the color that most closely represents their synesthetic sensation for a given letter or number. Letter presentation is repeated three times in random order. The variance for the three corresponding trials is calculated and serves as a means for synesthesia (for further explanation, see Eagleman et al., 2007). Two of the participants showed a variance score above the cut-off set by Eagleman (v ≤ 1), but reported to have difficulties finding the shimmering synesthetic colors in the test patch and, thus, were still classified as synesthetes. The mean variance score across the synesthesia group was 0.69 (range: 0.36 – 1.52). Synesthetes differ in the subjective quality of their color sensations. While ‘projector’ synesthetes perceive colors in the outside world on top of the printed graphemes, ‘associator’ synesthetes experience the colors only ‘in the mind’s eye’ (van Leeuwen, 2013). A modified german version of the projector-associator (PA) questionnaire introduced in Rouw and Scholte (2007) was used for assessing the synesthesia subtype (Volberg, Chockley, & Greenlee, 2017). Higher positive / negative scores indicate stronger projector / associator synesthesia, respectively. Four of the synesthetes were classified as associators while the other seven were classified as projectors.

Control participants did not complete the test battery but reported to not have any associations between colors and letters. They were recruited among the staff of the Psychology Department as well as from the pool of psychology students.

Compensation for participation was given either monetary (7-10 € per hour) or in the form of experimental hours for psychology students. All participants signed an informed consent form prior to the experiment.

### 2.2 MRI data acquisition

Brain images were acquired by a Siemens 3T Prisma MRI scanner with a 64 channel head coil (Erlangen, Germany). For the structural T1 weighted images, a magnetization-prepared rapid-acquisition gradient-echo sequence (MP-RAGE) across 160 sagittal slices and a voxel size of 1mm^3^ was used (TR = 2250 ms, TE = 2.6 ms, FA = 9°, TI = 900 ms, FOV = 256 × 256 mm^2^). The DWI were acquired with a single-shot spin-echo echo-planar sequence with a b-value of 1000 s/mm^2^ probing 30 isotropically distributed orientations across 60 axial slices with a voxel size of 2 × 2 × 2 mm^3^ (TR = 7700 ms, TE = 80 ms). Seven b-zero volumes were interspersed and an additional diffusion-weighted volume set with a higher b-value (2000 s/mm^2^, not analyzed here) were acquired. For the functional runs (retinotopy, color localizer, word localizer), gradient-echo echo-planar imaging (EPI) sequences (TR = 2s, TE = 30 ms, FA = 90°, voxel size = 3× 3× 3mm^3^, FOV = 192 × 192 mm^2^) across 37 transverse slices were used. The first five (retinotopy) or six (localizers) volumes of each run were discarded prior to the analysis to ensure a steady transverse magnetization.

### 2.3 Cortical Reconstruction

Structural images were automatically reconstructed by Freesurfer version 5 (Martinos Center for Biomedical Imaging, Charlestown, MA; Fischl, 2012). The reconstruction followed procedures as described in previous studies (Beer, Plank, & Greenlee, 2011). In brief, T1-weighted images of individual brains were intensity normalized and segmented into cortical gray and white matter. Then, the boundary between gray and white matter was tessellated and automatically corrected for topologic inaccuracies. Finally, the cortical surface was deformed, inflated, and registered to a spherical atlas that preserves the individual folding patterns of sulci and gyri (Fischl, Sereno, & Dale, 1999).

### 2.4 ROI definition

Diffusion parameters were assessed for several regions-of-interest (ROI) for both color processing and letter processing areas. ROIs were defined using multiple methods: based on an anatomical atlas, retinotopic mapping, functional group analysis, individual peak activations as well as combinations thereof. Due to our extensive use of ROI definitions, only left hemispheric ROIs are reported here. The left hemisphere is known for its dominance in language processing (Gerrits, Van der Haegen, Brysbaert, & Vingerhoets, 2019; Scheppele, Evans, & Brown, 2018), including atypical language experiences (Spray, Beer, Bentall, Sluming, & Meyer, 2018). Activation in occipital and temporal regions due to grapheme-color synesthesia seems to focus on the left hemisphere, as well (Rich et al., 2006).

#### 2.4.1 Anatomical ROI definition

For an initial analysis, eight ROIs (V4, V8, VVC, PH, PIT, PHT, FST, and MST, see Supplementary Figure S1) were defined anatomically using the Glasser Atlas (Glasser et al., 2016). This atlas separates cortical areas based on various parameters such as cortical thickness, myelination, and functional contrasts. Temporo-occipital areas V4, V8 and VVC were picked as candidates for color processing areas. Glassers V8 seems to be similar to the color sensitive area V8 of (Hadjikhani, Liu, Dale, Cavanagh, & Tootell, 1998), but also shows some overlap with VO1 (Abdollahi et al., 2014), an area found to be color selective (Brouwer & Heeger, 2009). VO1 is also included in Glassers VVC.

The VWFA is a functionally described area in the left fusiform gyrus (Cohen & Dehaene, 2004; Cohen et al., 2000). The Talairach coordinates reported for VWFA in the literature (Cohen & Dehaene, 2004; Cohen et al., 2000) overlap with area PH of Glasser et al. (2016). However, grapheme sensitive areas are likely located posterior to the VWFA (Thesen et al., 2012). Thus, we also included atlas-based areas located posterior but adjacent to it: PIT, PHT, FST and MST.

The eight anatomical areas were transformed to the participants’ individual cortical surfaces by Freesurfer based on the spherical registration. Surface areas were then converted into volumetric ROIs by projecting from the white-gray matter boundary 0 to 2 mm into the white matter.

#### 2.4.2 Functional ROI definition

Additional ROIs (V4r, ColorSphere, ColorFDR, WordSphere, WordFDR) were identified by task-based functional MRI activity patterns. All localizer tasks were run on a custom PC under the Psychophysics Toolbox for Matlab (Brainard, 1997; The MathWorks, Natick, MA, USA). The stimuli were projected on a translucent screen positioned about 95 cm distant to the eye, using a ProPixx VPXPRO5050A projector at a resolution of 1024×768 pixels and a vertical refresh rate of 60 Hz (VPixx Technologies, Inc., Quebec, Canada). Participants viewed the stimuli via a back-mirror that was mounted within the head-coil. MR-compatible button boxes were used as response devices.

##### 2.4.2.1 Retinotopic mapping

Color processing area V4 was retinotopically defined using the paradigm and analysis streamline of Frank, Reavis, Greenlee and Tse (2016). For better distinction from Glassers V4 within this article, the retinotopic area will be called V4r. The phase-encoded retinotopic mapping was conducted from two runs of a flickering colored bowtie-shaped double-wedge checkerboard rotating in clockwise (first run) and counterclockwise (second run) direction. Rotation traversed 18 screen positions for 3 seconds each for a total of 12 cycles. To ensure central fixation, participants engaged in a simple detection task. Analysis of the phase-encoded retinotopic mapping data followed standard techniques (Frank et al., 2016; Beer, Watanabe, Ni, Sasaki, & Andersen, 2009; DeYoe et al., 1996; Engel, Glover, & Wandell, 1997). For one synesthete, no retinotopic mapping could be performed due to technical problems during the measurement. Left V8 could only be identified in 8 of the 22 participants and, therefore, is not included in the further analysis.

##### 2.4.2.2 Color Localizer Task

Similar to the color localizer used by Gould van Praag, Garfinkel, Ward, Bor and Seth (2016) and Rich et al. (2006), the stimulation in this study consisted of alternating blocks of colored and grayscale patterns. Stimuli were constructed from 100 rectangles that were placed at random positions within a central screen area subtending 12 by 12 degrees of visual angle, on a uniformly black background. The rectangles had random edge lengths from 1 −2.5 degrees and were filled with random colors drawn from an RGB color space. Luminance-matched grayscale values were computed for the achromatic condition. Each block contained 24 different stimuli that were shown for 0.4s, with an 0.1s intermittent black screen. Color and grayscale blocks were separated by blocks of the same duration (12 s) where a blank screen was shown. Eight blocks with stimuli (four color, four grayscale) and eight blocks with a blank screen were show in one run, and three runs were administered overall. To ensure central fixation, the participants were instructed to to press a button whenever the central fixation blinked, using the index or middle finger of the same hand. A blink was 32 ms long, and the number of blinks was selected randomly (between 14 and 20) in each run.

##### 2.4.2.3 Word Localizer Task

Word sensitive areas were identified by a word localizer task similar to those described in the literature (Rauschecker, Bowen, Parvizi, & Wandell, 2012; Yeatman, Rauschecker, & Wandell, 2013). The word localizer task consisted of alternating blocks of german words, non-words, scrambled words and scrambled non-words, interleaved with 12s blank screen presentations. Common german words with four or five letters were chosen for the word condition, e.g. ‘park’ (park) or ‘gast’ (guest). Non-words were created by changing single letters and / or the order of the letters of the words, e. g. ‘karp’ or ‘stag’. The non-words didn’t contain semantic information in the german language but abide to linguistic rules. Words and non-words were written in 40 pt lowercase Helvetica font type, in white on black background. The text was converted into bitmap images where the size of the bitmap corresponded to the bounding box of the text (average height 1.4 degrees of visual angle). Scrambled versions of the images were constructed by computing Fourier transformations on the bitmap intensity and adding random values to the phase spectrum. The resulting images had the same average luminance as the original bitmaps, but don’t contain pictural information. All stimuli were presented in the screen center.

The trial sequence was identical to that of the color localizer task (24 stimuli per block, 0.4s presentation time plus 0.1s inter-stimulus interval). Two blocks per condition and eight blocks with a blank screen were shown in each run, and three runs were conducted overall. The task was to fixate the screen center and to indicate, by button press, a blink of the fixation spot.

##### 2.4.2.4 Analysis of the Localizers

All functional MRI data was analyzed by the FsFast tools of Freesurfer. Preprocessing included motion correction, skull stripping, intensity normalization and smoothing (full-width-half-maximum: 5mm) for each task separately. Functional runs were automatically linearly coregistered to individual anatomical space. All registrations were manually checked and corrected if necessary.

Both tasks, the color localizer and word localizer, were analyzed by a general linear model. First, an individual analysis was performed with the main contrasts ‘colored vs. gray patterns’ (color localizer) and ‘words and non-words versus scrambled’ (word localizer). Then, a surface-based group level random-effects analysis across all participants was performed to find a common cluster for the brain areas involved in color / word processing. For the color task, activation was found in the occipital, parietal and temporal cortex on both hemispheres. For the word localizer, activation was found in lateral occipital, lingual and middle temporal areas, as well as in the frontal lobe. According to the predictions of the cross-activation theory, we focused on occipito-temporal regions only. Supplementary Figure S2 shows the group clusters corrected for multiple comparisons with a voxelwise threshold of p = .001 and a cluster-wise threshold of p = .01.

The activated areas for color processing and word processing from the group analysis were transformed back to each subjects individual space by spherical registration. Similar to the approach by Rich et al. (2006) and Gould van Praag et al. (2016), a 10mm sphere was drawn around each participants peak activation to define their individual ColorSphere and WordSphere ROI, respectively. From these spheres, only those voxels that overlapped with the white matter of the cortical reconstruction and the group analysis cluster were kept (Supplementary Figure S3 A&C).

In addition, an alternative approach for the ROI definition based on individual activation patterns was used: Here, functional activation maps in each participant were FDR corrected (Benjamini & Hochberg, 1995; Yekutieli & Benjamini, 1999). For each participant, the functional ROIs for color and words were created from the individual significant voxels at an FDR adjusted p-value of 0.05. For the word localizer, no significant voxels survived the FDR correction in one synesthete. For the color localizer, no functional ROI could be extracted due to this problem in 2 synesthetes and 3 controls, limiting the further analysis of diffusion parameters within this area to groups of n=9 participants each. As with the other ROIs, from these ColorFDR and WordFDR ROIs only voxels overlapping with the white matter of the cortical reconstruction were kept (Supplementary Figure S3 B&D).

### 2.5 DTI Analysis

Diffusion data was processed by the FDT toolbox of FSL 5 (Oxford Center for Functional Magnetic Resonance Imaging of the Brain; Jenkinson, Beckmann, Behrens, Woolrich, & Smith, 2012). Diffusion images were corrected for head motion and eddy current distortions using the mid b-zero as reference. The b-zero volume was then linearly (rigid-body) co-registered to the individual structural image. Co-registrations were manually checked and corrected if necessary. Diffusion parameters fractional anisotropy (FA), mean diffusivity (MD), axial diffusivity (AD) and radial diffusivity (RD) were estimated by the diffusion tensor model (Basser, Mattiello, & LeBihan, 1994). Then, mean diffusion parameters were calculated for each ROI first for each individual and then across groups.

### 2.6 Statistical analysis

Statistics were computed for group differences in mean diffusion parameter values, FA and MD, between synesthetes and controls. This was done separately for each ROI.

Individual diffusion parameters were entered into a Bayesian t-test for independent groups. If synesthetes were to exhibit increased structural connectivity, their FA values should be higher than the values of the control group, and their MD values should be lower, compared to the control group. That is, the true effect size *δ* should be positive for FA (H_1_: δ ≥ 0, H_0_: *δ* = 0) and negative for MD as dependent variables (H_1_: *δ* ≤ 0, H_0_: *δ* = 0). Other than in conventional significance testing, the evidence for a null hypothesis can be assessed without knowing the true effect size by imposing constraints on the effect size distribution. Cauchy priors with scale parameter r = √(2)/2 were used (Morey & Rouder, 2011b). About 40% of effect sizes in this distribution are within the interval from −0.5 to 0.5, reflecting small-to-medium effects. For comparison, we also report the results of Welch’s t-test for independent samples with unequal variances. One-sided tests were used, analogous to the computation of Bayesian factors (H_1_for FA: μ_synesthetes_-μ_controls_. > 0; for MD: μ_synesthetes_-μ_controls_. < 0). T-tests were not corrected for multiple comparisons, in order to maximize the statistical power.

Previous research suggests that structural brain differences are more likely to occur in the subgroup of projector synesthetes with perception-like color sensations for graphemes, compared to associator synesthetes (van Leeuwen et al., 2011). In addition to group differences in DTI parameters between synesthetes and controls, possible association between PA scores and diffusion parameters were assessed within the group of synesthetes. Pearson correlations with the PA score were computed per diffusion metric and ROI. Average correlations for color and grapheme ROIs were obtained by transforming r into z_r_, averaging z_r_, and back-transforming the mean z_r_into r using Fisher’s z formula (Fisher, 1915).

Negative PA scores are indicative for the associator type and positive scores indicate the projector type of synesthesia. Given that increased neural cross-wiring is reflected in high FA values and low MD values, and assuming that cross-wiring is more prominent in projector synesthetes, the analysis should reveal positive correlations between FA and PA values and negative correlations between MD and PA scores. Bayesian factors were obtained for the hypothesis that the true linear correlation `ρ` between PA score and FA values is larger than zero (H_1_: `ρ` ≥ 0; H_0_: `ρ` = 0), and for the hypothesis that the true correlation is smaller than zero (H_1_: `ρ` ≤ 0; H_0_: `ρ` = 0). Beta distributions with scale 1/3 were used as priors.

The R package for statistical computing (R Core Team, 2018) and the package ‘BayesFactor’ (Rouder, Speckman, Sun, Morey, & Iverson, 2009) were used for conducting the statistics and for plotting the results.

## 3. Results

### 3.1. DTI parameters: Group differences

#### 3.1.1 Color processing areas

We investigated diffusion parameters FA and MD as means of coherent fibre tracts and myelination in three anatomically and three functionally defined regions of interest for color processing. Results are depicted in Figure 1 and Figure 3, the details of the statistical analysis are given in supplementary Tables S1 (FA values) and S2 (MD values).

**Figure 1.**
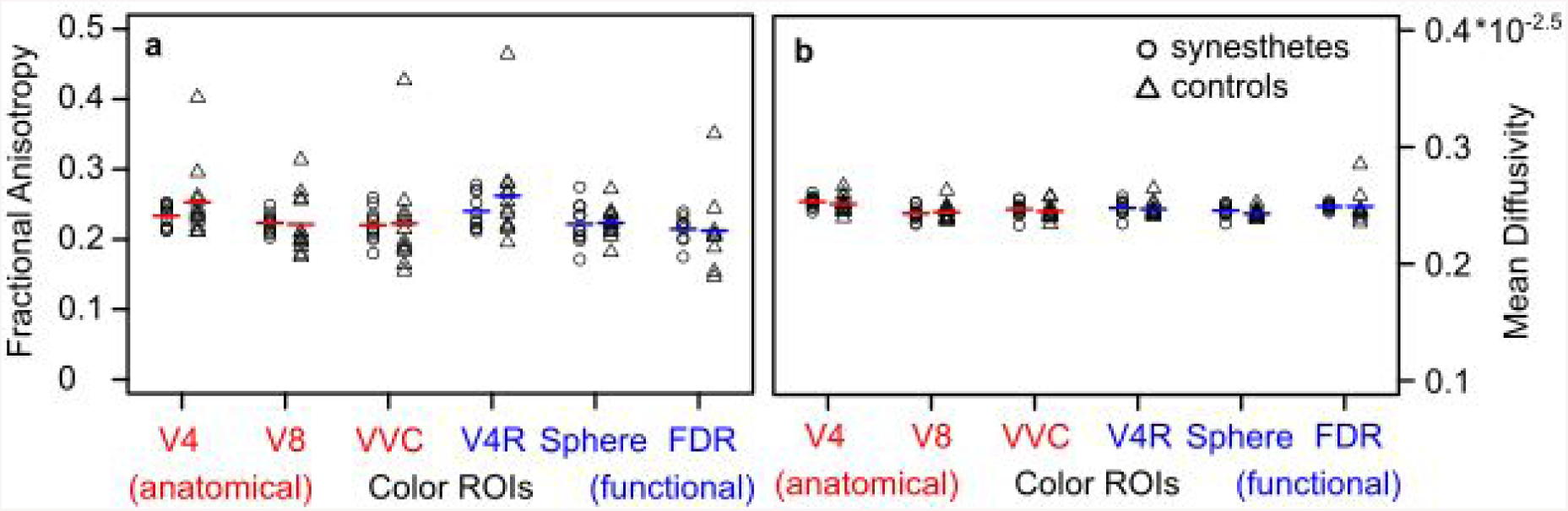
Diffusion parameters obtained from color ROIs, contingent on participant groups. Mind the different scales for fractional anisotropy (a) and mean diffusivity data (b). Vertical bars indicate the arithmetic mean. See text for ROI definitions.

Mean FA values were somewhat larger in controls compared to synesthetes in anatomically and functionally defined V4, but had similar sizes for both groups in the remaining ROIs. The mean BF_10_was 0.33 (range 0.21 −0.44). Thus, the hypothesis of a null difference of FA values between groups was 3.25 times (range 2.28 −4.85) more likely than the H1, given the data. Correspondingly, Welch’s t-tests did not reach significance in any case (all t ≤ 0.144, all p > .44).

MD values were slightly lower, on average, for synesthetes compared to controls in areas V8 and C-FDR, and slightly higher in all other ROIs. The mean BF_10_was 0.32 (0.19 − 0.52). That is, the null hypothesis was 3.49 (1.94 − 5.36) times more likely than the experimental hypothesis given the data. None of the t-tests on group differences in MD values was significant, all t ≥ −0.427, all p > .33.

Visual inspection of Figure 1 shows that the variance of FA values (not so in MD values) was often higher in controls compared to synesthetes. This was caused by one participant with untypically high FA values. Outlier values could inflate the group averages for controls and render the expected differences to synesthetes undetectable. In order to ensure that outliers did not affect the obtained results, the FA-value analysis was repeated after removing data of the untypical control participant. The overall results pattern was similar to that of the previous analysis. BFs favored the alternative hypothesis for region VVC, but the null hypothesis in all other ROIs. The mean BF_10_was 0.61 (0.26 − 1.33), indicating that the H_0_was 2.20 (0.75 − 3.77) times more likely than the H_1_. Welch’s t-tests were insignificant, all t < 1.40, p > .089.

#### 3.1.2 Letter processing areas

FA and MD were investigated in five anatomically and two functionally defined regions of interest for letter processing. The results are summarized in Figure 2 and Figure 3. Details of the statistical analysis are given in supplementary Tables S3 (FA values) and S4 (MD values).

**Figure 2.**
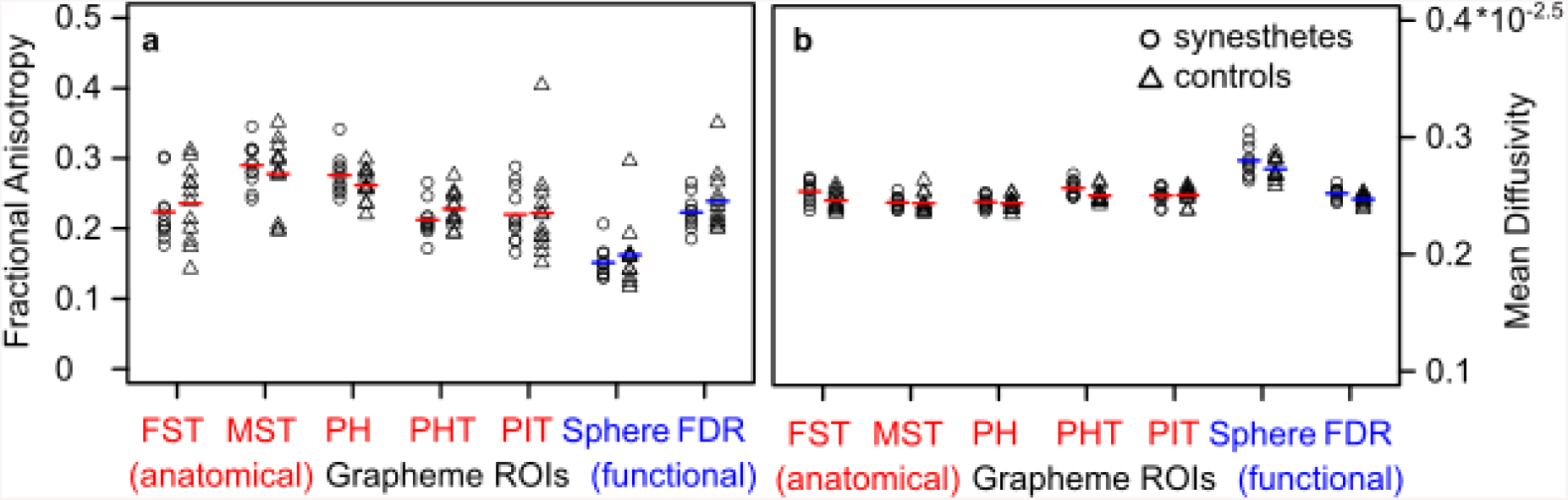
Same as Figure 1, but for diffusion parameters obtained from grapheme processing ROIs.

**Figure 3.**
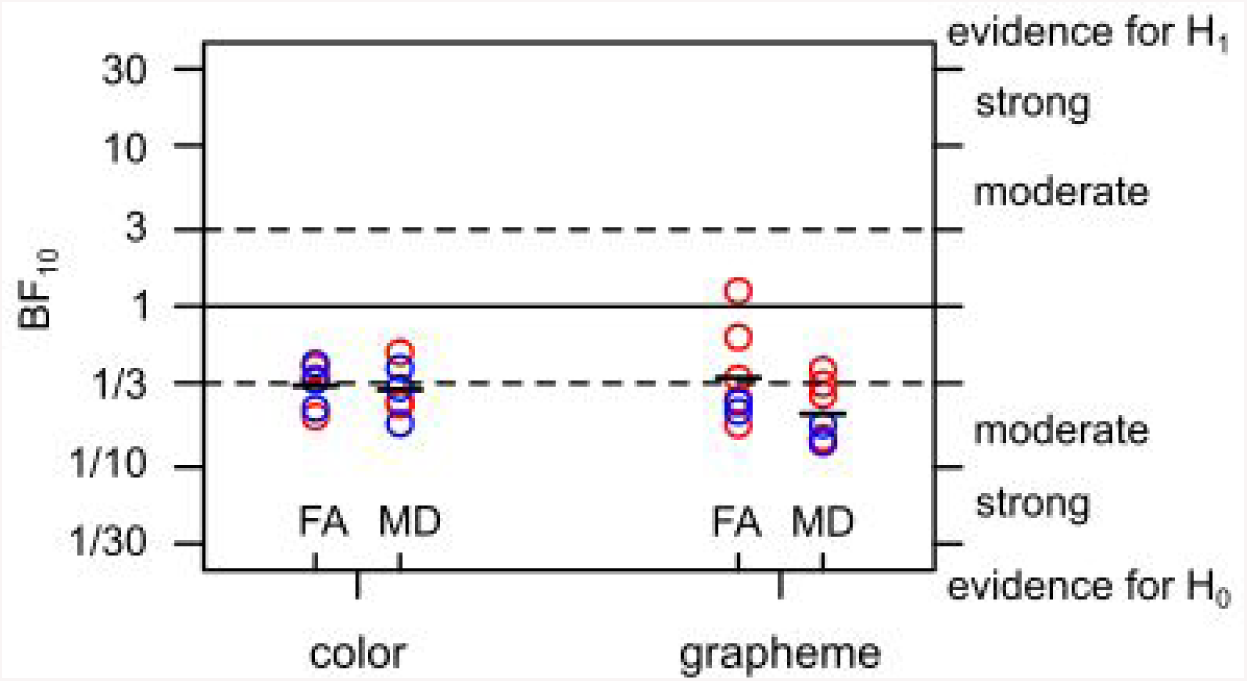
Bayesian inference on group differences between synesthetes and controls. Each circle represents the Bayesian factor (BF) for one anatomically (red) or functionally (blue) defined color or grapheme ROI. Vertical bars indicate the mean BF_10_across color and grapheme ROIs, respectively. Values < 1 support the hypothesis of a null difference between groups. Dashed lines mark BF values that are considered a moderate effect.

The mean FA values were somewhat higher in synesthetes than in controls in areas MST and PH, and lower in all other regions. BFs_10_favored the alternative hypothesis over the null hypothesis in ROI PH, but not in the other ROIs. On average, the hypothesis of a null difference in FA values was 3.26 (0.81 − 5.44) times more likely than the alternative hypothesis of a group difference. Welch’s t-tests were insignificant for all ROIs, all t ≤ 1.33, all p > .099.

With respect to the MD parameter, synesthetes showed slightly lower mean values than controls in area PIT, and higher values in the other ROIs. Bayesian testing favored the H_0_over the H_1_in all ROIs (mean BF_10_= 0.23, range 0.14 − 0.40; or BF_01_= 5.02, 2.49 − 7.06). All t-tests were insignificant, t ≥ -.088, p > .46.

### 3.2 DTI parameters in synesthetes: Correlations

Pearson correlations between PA scores and FA or MD diffusion parameters were calculated per ROI. The results are depicted in Figure 4. Dashed lines show the corresponding regression slope for each ROI, the solid black bar marks the average correlation. Bayesian factors were calculated for the H1 that the correlation differs from 0, versus the H0 of a null correlation.

**Figure 4.**
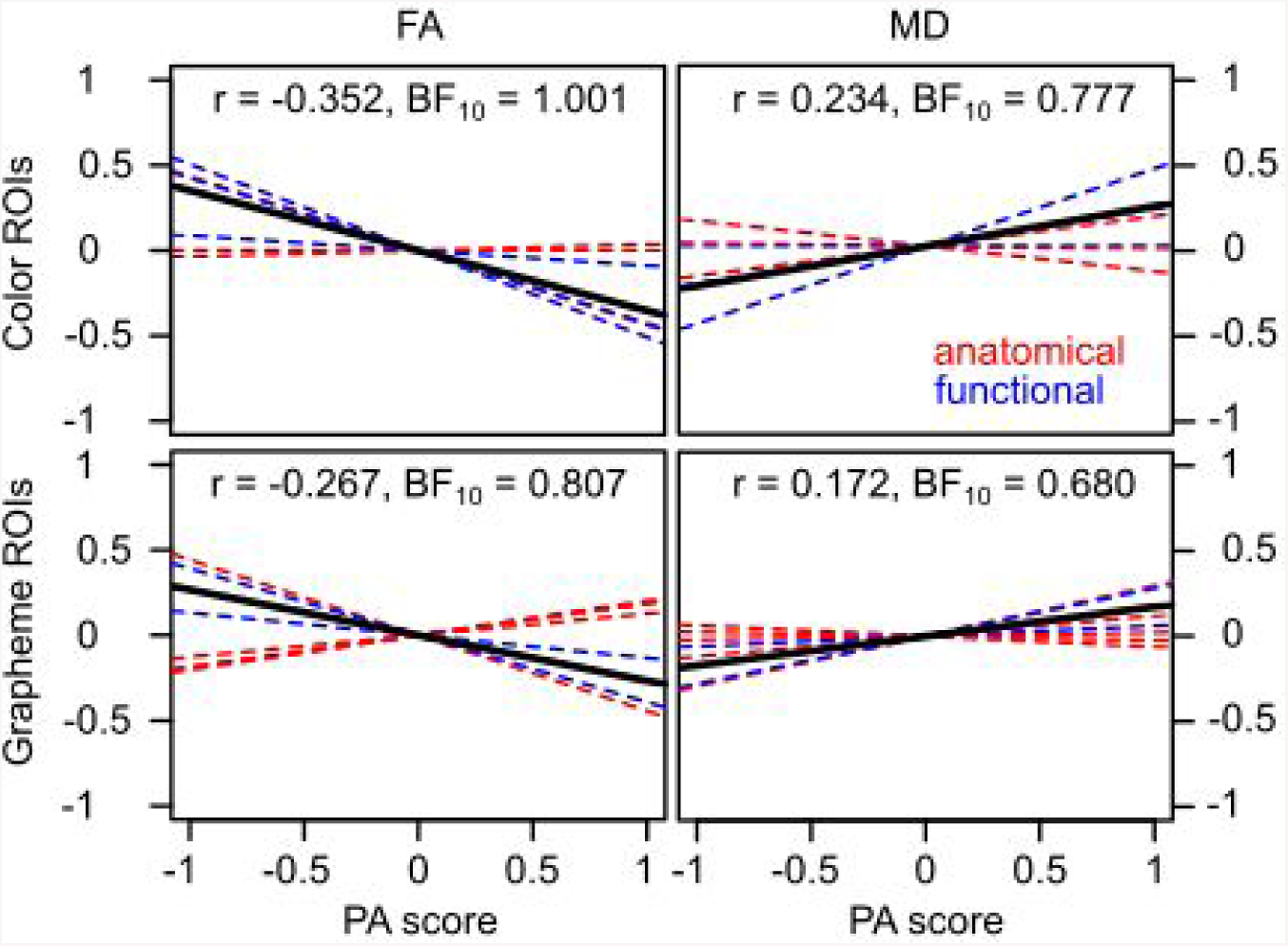
Correlation between projector-associator scores (PA) and diffusion parameters ROIs in the group of synesthetes. Dashed lines mark slopes for correlations obtained in single anatomically (red) or functionally (blue) defined color and grapheme ROIs. Black solid lines mark the average correlation per diffusion parameter and ROI group.

If structural connectivity is stronger in projector synesthetes with higher scores in the PA scale, then positive correlations should occur between PA and FA parameters (where high values reflect strong connectivity). Such a pattern was observed in only one out of six color ROIs. Correlations ranged from r = −0.51 to r = 0.04, the average correlation was −0.352. With respect to letter ROIs, correlations were positive in four out of seven cases. Nonetheless, the average correlation was negative, r = −0.267, range r = −0.44 to r = 0.19.

With respect to MD parameters, where low values reflect high structural connectivity, negative correlations with PA scores should be observed. Thus was the case in only two out of six color ROIs, range r = −0.15 to r = 0.45, mean r = 0.234, and in two out of seven grapheme ROIs, range r = −0.06 to 0.30, mean r = 0.172.

The BF_10_ were generally inconclusive. Overall, the Bayesian factors favored the hypothesis of a null correlation between PA scores and FA/MD values, which was 0.99 − 1.47 times more likely than the alternative hypothesis given the data.

## 4. Discussion

The aim of this study was to investigate possible microstructural differences between synesthetes and non-synesthetes in brain areas proposed by the cross-activation theory. This theory proposes a hyperconnectivity leading to immediate cross-talk between grapheme processing areas, likely VWFA, and color processing areas, likely V4, in grapheme-color synesthetes (Hubbard et al., 2011; Ramachandran & Hubbard, 2001). To test this, we examined diffusion parameters FA and MD as measures of white matter coherence in 13 ROIs, six color sensitive areas and seven candidates for the VWFA, in a sample of 11 synesthetes and 12 non-synesthetic controls. To counter possible null effects due to inaccurate localization of our investigations, multiple methods were used to define the ROIs anatomically and functionally. Nevertheless, none of our ROIs revealed sufficient group differences to support the assumptions of the cross-activation theory.

The most prominent color-sensitive area V4 was defined atlas-based as well as with retinotopic mapping. In both anatomical and retinotopic V4, FA was slightly increased for synesthetes compared to controls (see Fig.1). However, this difference was not significant. Furthermore, atlas-based areas V8 and VVC (including VO1) were examined. Using a functional color localizer and two different analysis techniques, two functional color-sensitive ROIs were defined. None of the color ROIs showed a group difference in FA values. In addition to the results obtained by standard significance testing, the results of Bayesian testing support the hypothesis of a null difference between the groups. The same conclusion can be drawn for MD within the color ROIs. Hence, our data suggests that there are no relevant microstructural differences between synesthetes and non-synesthetes in perceptual color processing areas. This finding is in line with previous studies, that did not find any structural differences in V4 or locations nearby (Dojat et al., 2018; Hupé & Dojat, 2015; Rouw & Scholte, 2007).

Since the VWFA is a functional and not anatomically specified area, we examined five atlas-based ROIs surrounding the Talairach coordinates reported in fMRI studies on VWFA. Two functional ROIs were defined using a word localizer task. Synesthetes showed higher FA values only in two anatomical ROIs, MST and PH (see Fig.2). The group differences did not reach significance in any grapheme ROI, though. Bayesian testing favors the null hypothesis except for area PH. Since PH shows the most overlap with VWFA related Talairach coordinates, this region might be the best candidate for an atlas-based alternative to the functional VWFA. While further analysis of this area might be promising, the FA values found in the present study were not high enough in synesthetes to produce a significant group difference. For MD, no significant group differences could be detected and Bayesian testing likewise supported the null hypothesis of no group differences. Similar results were found by (Jäncke et al., 2009), who did not detect significant group differences of FA in the fusiform gyrus. They proposed that structural anomalies in synesthetes might be located around V4. However, our findings suggest that neither processing region, V4 nor VWFA, exhibits increased white matter coherence in synesthetes.

Other studies have been criticized for their use of atlas-based ROI definition (Banissy et al., 2012; Weiss & Fink, 2009). Since this study used multiple definitions of the ROIs, a direct comparison between anatomical and functional definitions can be drawn. For the color processing areas, the three atlas-based (V4, V8, VVC) and the three functionally defined (V4r, ColorSphere, ColorFDR) areas hold FA and MD values in the same range. In all six ROIs, the null hypothesis of no group differences for FA and MD is supported by moderate Bayesian factor strengths. The VWFA is an originally functionally defined area. We used a similar functional localizer as stated in the literature (Yeatman et al., 2013). The ROI emanating from the individually FDR adjusted significant voxels for words and non-words vs scrambled words (WordFDR) contains similar MD values to the atlas-based ROIs. FA values differed slightly between the seven ROIs but in no particular direction. Significance testing revealed no group differences in any of the ROIs. Bayesian testing also showed no pattern of better fit for either atlas-based or functional ROIs. Thus, none of our methods would improve the detection of group differences between synesthetes and controls in regards to the diffusion parameters.

It has been argued that a structural hyperconnectivity is found preferentially in projector type synesthetes (Rouw & Scholte, 2007, 2010). Their synesthesia is expressed in atypical perception of the outer world and, therefore, might rely more on neural activity within perceptual processing areas. Dense axonal packing and high myelination should be exhibited in higher FA and lower MD values in these subjects compared to associator type synesthetes. Contrary to that prediction, no correlation could be found between the FA and MD values and type of synesthesia. Correlations were generally weak and insignificant, and the Bayesian factor values were low. Due to the small sample size, the statistics are only tentative. However, it is safe to say that our data do not show altered microstructure, even in a selection of projector synesthetes. It should be mentioned that the distinction between projector and associator synesthetes is considered inappropriate by some authors (Hupé & Dojat, 2015). Indeed, some of the synesthetes investigated in this study had difficulties describing their individual perception and gave contradicting answers in the PA-questionnaire. It might be hard to find clearly defined groups of different types of grapheme-color synesthetes and to delineate brain differences between them.

Taken together, we examined diffusion parameters in functionally and anatomically defined color- and grapheme-sensitive ROIs, from individual cortical sheets, and found no group differences in FA or MD. Also, there were no differences between synesthetes with perception-like (projectors) or memory-like (associators) color experiences. Bayesian statistics revealed moderate-to-strong support across ROIs for the null hypothesis of no group differences in white matter microstructure in the brain of synesthetes and controls. The findings argue against a cross-activation of color as the source of the synesthetic perception.

## Acknowledgements

This work was supported by the Deutsche Forschungsgemeinschaft, VO1998/1-1. We thank Anton Beer, Markus Becker and Anna Saga Wilhelmina Lee Örbom for their contribution.

---

**Table S1.** Statistics for fractional anisotropy (FA) in the color ROIs.

| ROI | Mean ± SD | | t | df | p | B<sub>10</sub> (BF<sub>01</sub>) |
| --- | --- | --- | --- | --- | --- | --- |
|  | synesthetes | controls |  |  |  |  |
| Glasser V4 | 0.233 ± 0.016 | 0.253 ± 0.053 | -1.210 | 13.1 | .877 | 0.206 (4.848) |
| Glasser V8 | 0.223 ± 0.014 | 0.221 ± 0.043 | 0.144 | 13.5 | .444 | 0.417 (2.398) |
| Glasser VVC | 0.220 ± 0.023 | 0.222 ± 0.071 | -0.092 | 13.5 | .536 | 0.358 (2.796) |
| V4R | 0.240 ± 0.025 | 0.262 ± 0.069 | -1.010 | 14.4 | .836 | 0.232 (4.318) |
| Sphere | 0.222 ± 0.029 | 0.223 ± 0.022 | -0.135 | 18.7 | .553 | 0.347 (2.882) |
| FDR | 0.215 ± 0.020 | 0.212 ± 0.060 | 0.099 | 9.71 | .462 | 0.440 (2.275) |

**Table S2.** Statistics for mean diffusivity (MD) in the color ROIs.

| ROI | Mean ± SD [$\times 10^{-2.5}$] | | t | df | p | $B_{10}$ ($BF_{01}$) |
| --- | --- | --- | --- | --- | --- | --- |
|  | synesthetes | controls |  |  |  |  |
| Glasser V4 | 0.253 ± 0.005 | 0.251 ± 0.007 | 0.797 | 18.6 | 0.782 | 0.244 (4.102) |
| Glasser V8 | 0.243 ± 0.006 | 0.244 ± 0.007 | -0.427 | 20.7 | 0.337 | 0.516 (1.938) |
| Glasser VVC | 0.247 ± 0.006 | 0.245 ± 0.007 | 0.670 | 21.0 | 0.745 | 0.258 (3.879) |
| V4R | 0.248 ± 0.006 | 0.247 ± 0.007 | 0.352 | 19.6 | 0.636 | 0.311 (3.216) |
| Sphere | 0.246 ± 0.005 | 0.243 ± 0.004 | 1.42 | 18.9 | 0.914 | 0.186 (5.364) |
| FDR | 0.249 ± 0.003 | 0.249 ± 0.015 | -0.001 | 8.55 | 0.497 | 0.440 (2.419) |

**Table S3.** Statistics for fractional anisotropy (FA) in the grapheme ROIs.

| ROI | Mean ± SD | | t | df | p | **B<sub>10</sub> (BF<sub>01</sub>)** |
| --- | --- | --- | --- | --- | --- | --- |
|  | synesthetes | controls |  |  |  |  |
| Glasser PH | 0.276 ± 0.028 | 0.262 ± 0.023 | 1.33 | 19.5 | .099 | 1.25 (0.802) |
| Glasser MST | 0.290 ± 0.030 | 0.278 ± 0.052 | 0.698 | 18.0 | .247 | 0.642 (1.558) |
| Glasser FST | 0.224 ± 0.042 | 0.236 ± 0.054 | -0.621 | 20.5 | .729 | 0.265 (3.778) |
| PHT | 0.212 ± 0.025 | 0.228 ± 0.025 | -1.48 | 20.9 | .923 | 0.184 (5.437) |
| PIT | 0.220 ± 0.040 | 0.222 ± 0.066 | -0.095 | 18.3 | .537 | 0.357 (2.804) |
| Sphere | 0.152 ± 0.022 | 0.163 ± 0.047 | -0.727 | 15.9 | .761 | 0.253 (3.957) |
| FDR | 0.223 ± 0.024 | 0.239 ± 0.043 | -1.090 | 17.9 | .855 | 0.222 (4.508) |

**Table S4.** Statistics for mean diffusivity (MD) in the grapheme ROIs.

| ROI | Mean ± SD [$\times 10^{-2.5}$] | | t | df | p | B10 (BF01) |
| --- | --- | --- | --- | --- | --- | --- |
|  | synesthetes | controls |  |  |  |  |
| Glasser PH | 0.245 ± 0.005 | 0.243 ± 0.006 | 0.503 | 20.9 | .690 | 0.280 (3.566) |
| Glasser MST | 0.254 ± 0.005 | 0.246 ± 0.009 | 2.11 | 19.9 | .589 | 0.329 (3.039) |
| Glasser FST | 0.244 ± 0.009 | 0.244 ± 0.008 | 0.227 | 18.2 | .976 | 0.152 (6.598) |
| PHT | 0.256 ± 0.007 | 0.250 ± 0.006 | 2.40 | 20.6 | .987 | 0.142 (7.057) |
| PIT | 0.250 ± 0.007 | 0.250 ± 0.007 | -0.088 | 20.7 | .466 | 0.402 (2.486) |
| Sphere | 0.280 ± 0.013 | 0.273 ± 0.009 | 1.47 | 17.5 | .920 | 0.183 (5.466) |
| FDR | 0.252 ± 0.005 | 0.247 ± 0.004 | 2.42 | 17.9 | .987 | 0.144 (6.943) |

**Figure S1.**
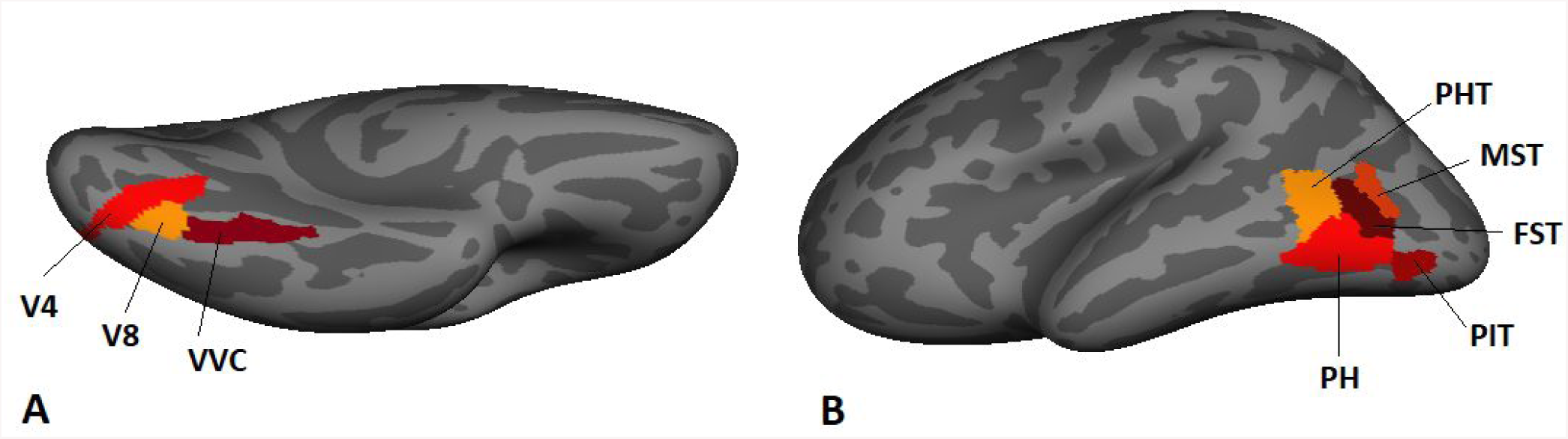
Anatomical labels for color selective areas (A) and possible word processing areas (B) from the Glasser Atlas on the left hemisphere of the freesurfer fsaverage

**Figure S2.**
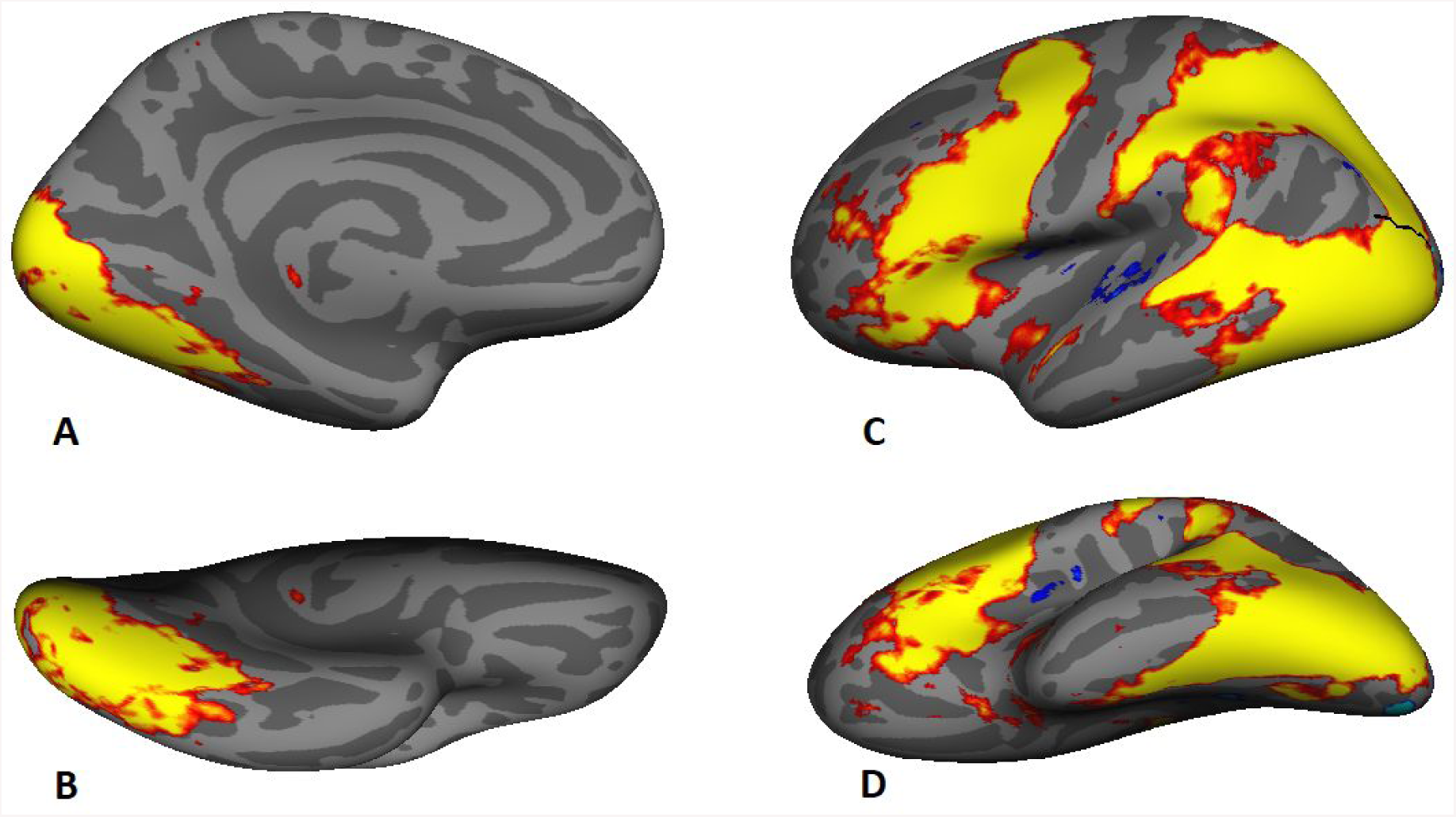
Significant clusters of the second level group analysis for the color localizer (contrast colored vs gray patterns, A & B), and the word localizer (contrast words and non-words vs scrambled, C & D) on the left hemisphere of the freesurfer fsaverage

**Figure S3.**
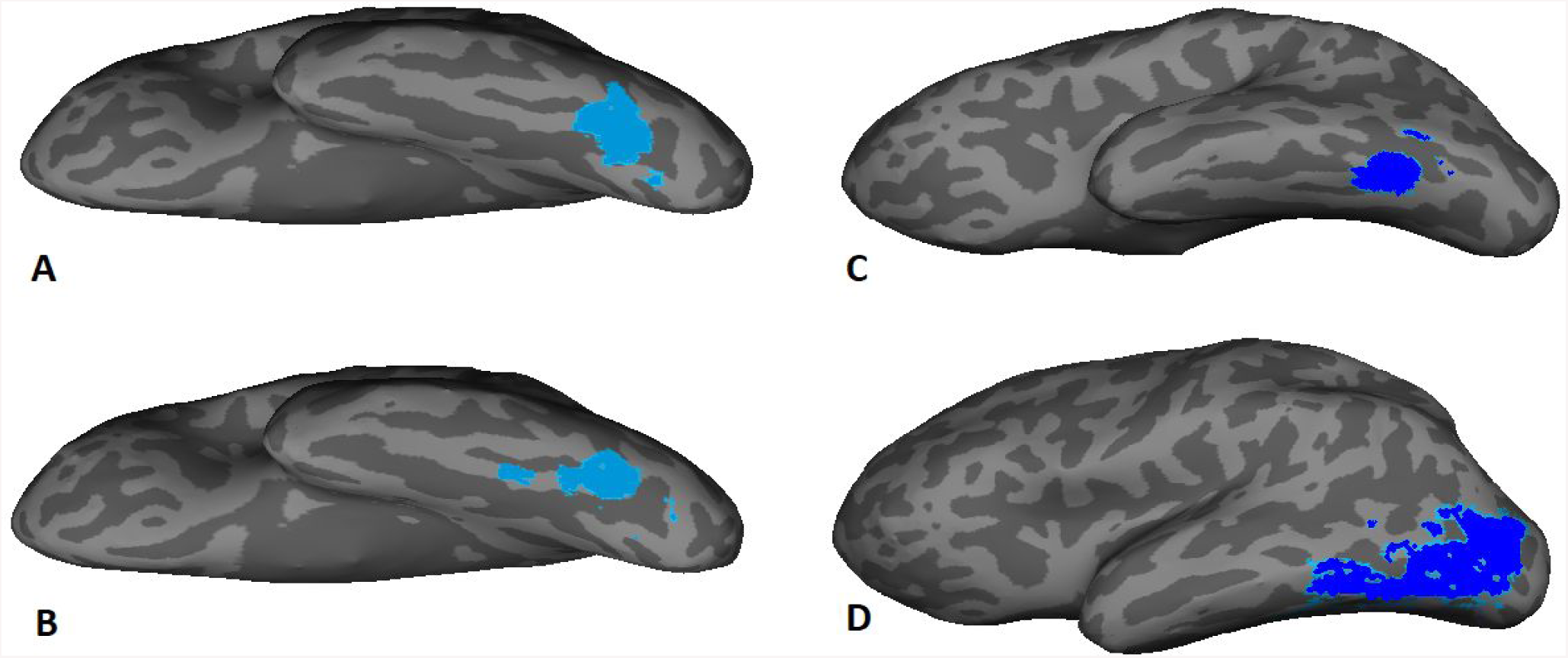
Functional ROIs of one exemplary synesthete. A 10mm sphere was set around the individual peak activation and masked with the white matter portion of the corresponding group cluster (A: color localizer, C: word localizer). For a second analysis, the individual FDR adjusted significant voxels within the group cluster were kept as ROI for color (B) and word processing (D).

## References

Abdollahi, R. O., Kolster, H., Glasser, M. F., Robinson, E. C., Coalson, T. S., Dierker, D., Orban, G. A. (2014). Correspondences between retinotopic areas and myelin maps in human visual cortex. NeuroImage, 99, 509–524.

Arend, I., Yuen, K., Sagi, N., & Henik, A. (2018). Neuroanatomical basis of number synaesthesias: A voxel-based morphometry study. Cortex; a Journal Devoted to the Study of the Nervous System and Behavior, 101, 172–180.

Banissy, M. J., Stewart, L., Muggleton, N. G., Griffiths, T. D., Walsh, V. Y., Ward, J., & Kanai, R. (2012). Grapheme-color and tone-color synesthesia is associated with structural brain changes in visual regions implicated in color, form, and motion. Cognitive Neuroscience, 3(1), 29–35.

Bartolomeo, P., Bachoud-Lévi, A.-C., & Thiebaut de Schotten, M. (2014). The anatomy of cerebral achromatopsia: a reappraisal and comparison of two case reports. Cortex; a Journal Devoted to the Study of the Nervous System and Behavior, 56, 138–144.

Basser, P. J., Mattiello, J., & LeBihan, D. (1994). MR diffusion tensor spectroscopy and imaging. Biophysical Journal, 66(1), 259–267.

Beer, A. L., Plank, T., & Greenlee, M. W. (2011). Diffusion tensor imaging shows white matter tracts between human auditory and visual cortex. Experimental Brain Research. Experimentelle Hirnforschung. Experimentation Cerebrale, 213(2-3), 299–308.

Beer, A. L., Watanabe, T., Ni, R., Sasaki, Y., & Andersen, G. J. (2009). 3D surface perception from motion involves a temporal-parietal network. The European Journal of Neuroscience, 30(4), 703–713.

Benjamini, Y., & Hochberg, Y. (1995). Controlling the False Discovery Rate: A Practical and Powerful Approach to Multiple Testing. Journal of the Royal Statistical Society: Series B (Methodological). https://doi.org/10.1111/j.2517-6161.1995.tb02031.x

Brainard, D. H. (1997). The Psychophysics Toolbox. Spatial Vision, 10, 433–436.

Brouwer, G. J., & Heeger, D. J. (2009). Decoding and reconstructing color from responses in human visual cortex. The Journal of Neuroscience: The Official Journal of the Society for Neuroscience, 29(44), 13992–14003.

Cohen, L., & Dehaene, S. (2004). Specialization within the ventral stream: the case for the visual word form area. NeuroImage, 22(1), 466–476.

Cohen, L., Dehaene, S., Naccache, L., Lehéricy, S., Dehaene-Lambertz, G., Hénaff, M.-A., & Michel, F. (2000). The visual word form area. Brain. https://doi.org/10.1093/brain/123.2.291

Cohen, L., Dehaene, S., Naccache, L., Lehéricy, S., Dehaene-Lambertz, G., Hénaff, M. A., & Michel, F. (2000). The visual word form area: spatial and temporal characterization of an initial stage of reading in normal subjects and posterior split-brain patients. Brain: A Journal of Neurology, 123 (Pt 2), 291–307.

Coriat, I. H. (1913). A case of synesthesia. The Journal of Abnormal Psychology. https://doi.org/10.1037/h0072314

DeYoe, E. A., Carman, G. J., Bandettini, P., Glickman, S., Wieser, J., Cox, R., … Neitz, J. (1996). Mapping striate and extrastriate visual areas in human cerebral cortex. Proceedings of the National Academy of Sciences of the United States of America, 93(6), 2382–2386.

Dojat, M., Pizzagalli, F., & Hupe, J.-M. (2018). Magnetic resonance imaging does not reveal structural alterations in the brain of synesthetes. https://doi.org/10.1101/196865

Eagleman, D. M., Kagan, A. D., Nelson, S. S., Sagaram, D., & Sarma, A. K. (2007). A standardized test battery for the study of synesthesia. Journal of Neuroscience Methods, 159(1), 139–145.

Engel, S. A., Glover, G. H., & Wandell, B. A. (1997). Retinotopic organization in human visual cortex and the spatial precision of functional MRI. Cerebral Cortex, 7(2), 181–192.

Fischl, B. (2012). FreeSurfer. NeuroImage, 62(2), 774–781.

Fischl, B., Sereno, M. I., & Dale, A. M. (1999). Cortical surface-based analysis. II: Inflation, flattening, and a surface-based coordinate system. NeuroImage, 9(2), 195–207.

Fisher, R. A. (1915). Frequency Distribution of the Values of the Correlation Coefficient in Samples from an Indefinitely Large Population. Biometrika. https://doi.org/10.2307/2331838

Frank, S. M., Reavis, E. A., Greenlee, M. W., & Tse, P. U. (2016). Pretraining Cortical Thickness Predicts Subsequent Perceptual Learning Rate in a Visual Search Task. Cerebral Cortex, 26(3), 1211–1220.

Gerrits, R., Van der Haegen, L., Brysbaert, M., & Vingerhoets, G. (2019). Laterality for recognizing written words and faces in the fusiform gyrus covaries with language dominance. Cortex. https://doi.org/10.1016/j.cortex.2019.03.010

Glasser, M. F., Coalson, T. S., Robinson, E. C., Hacker, C. D., Harwell, J., Yacoub, E., … Van Essen, D. C. (2016). A multi-modal parcellation of human cerebral cortex. Nature, 536(7615), 171–178.

Gould van Praag, C. D., Garfinkel, S., Ward, J., Bor, D., & Seth, A. K. (2016). Automaticity and localisation of concurrents predicts colour area activity in grapheme-colour synaesthesia. Neuropsychologia, 88, 5–14.

Hadjikhani, N., Liu, A. K., Dale, A. M., Cavanagh, P., & Tootell, R. B. (1998). Retinotopy and color sensitivity in human visual cortical area V8. Nature Neuroscience, 1(3), 235–241.

Hubbard, E. M., Brang, D., & Ramachandran, V. S. (2011). The cross-activation theory at 10. Journal of Neuropsychology, 5(2), 152–177.

Hupé, J.-M., Bordier, C., & Dojat, M. (2012). The neural bases of grapheme-color synesthesia are not localized in real color-sensitive areas. Cerebral Cortex, 22(7), 1622–1633.

Hupé, J.-M., & Dojat, M. (2015). A critical review of the neuroimaging literature on synesthesia. Frontiers in Human Neuroscience, 9, 103.

Jäncke, L., Beeli, G., Eulig, C., & Hänggi, J. (2009). The neuroanatomy of grapheme-color synesthesia. The European Journal of Neuroscience, 29(6), 1287–1293.

Jenkinson, M., Beckmann, C. F., Behrens, T. E. J., Woolrich, M. W., & Smith, S. M. (2012). FSL. NeuroImage, 62(2), 782–790.

Lee, M. D., & Wagenmakers, E.-J. (2014). Bayesian Cognitive Modeling: A Practical Course. Cambridge University Press.

McCandliss, B. D., Cohen, L., & Dehaene, S. (2003). The visual word form area: expertise for reading in the fusiform gyrus. Trends in Cognitive Sciences, 7(7), 293–299.

McErlean, A. B. J., Janik McErlean, A. B., & Banissy, M. J. (2017). Color Processing in Synesthesia: What Synesthesia Can and Cannot Tell Us About Mechanisms of Color Processing. Topics in Cognitive Science. https://doi.org/10.1111/tops.12237

Morey, R. D., & Rouder, J. N. (2011a). Bayes factor approaches for testing interval null hypotheses. Psychological Methods, 16(4), 406–419.

Morey, R. D., & Rouder, J. N. (2011b). Bayes factor approaches for testing interval null hypotheses. Psychological Methods, 16(4), 406–419.

Ramachandran, V. S., & Hubbard, E. M. (2001). Psychophysical investigations into the neural basis of synaesthesia. Proceedings. Biological Sciences / The Royal Society, 268(1470), 979–983.

Rauschecker, A. M., Bowen, R. F., Parvizi, J., & Wandell, B. A. (2012). Position sensitivity in the visual word form area. Proceedings of the National Academy of Sciences of the United States of America, 109(24), E1568–E1577.

R Core Team. (2018). R: A Language and Environment for Statistical Computing. Vienna, Austria: R Foundation for Statistical Computing. Retrieved from https://www.R-project.org/

Rich, A. N., Williams, M. A., Puce, A., Syngeniotis, A., Howard, M. A., McGlone, F., & Mattingley, J. B. (2006). Neural correlates of imagined and synaesthetic colours. Neuropsychologia. https://doi.org/10.1016/j.neuropsychologia.2006.06.024

Roe, A. W., Chelazzi, L., Connor, C. E., Conway, B. R., Fujita, I., Gallant, J. L., … Vanduffel, W. (2012). Toward a Unified Theory of Visual Area V4. Neuron. https://doi.org/10.1016/j.neuron.2012.03.011

Rouder, J. N., Speckman, P. L., Sun, D., Morey, R. D., & Iverson, G. (2009). Bayesian t tests for accepting and rejecting the null hypothesis. Psychonomic Bulletin & Review, 16(2), 225–237.

Rouw, R., & Scholte, H. S. (2007). Increased structural connectivity in grapheme-color synesthesia. Nature Neuroscience, 10(6), 792–797.

Rouw, R., & Scholte, H. S. (2010). Neural basis of individual differences in synesthetic experiences. The Journal of Neuroscience: The Official Journal of the Society for Neuroscience, 30(18), 6205–6213.

Scheppele, M., Evans, J. L., & Brown, T. T. (2018). Patterns of structural lateralization in cortical language areas of older adolescents. Laterality, 1–32.

Smith, S. M., Jenkinson, M., Johansen-Berg, H., Rueckert, D., Nichols, T. E., Mackay, C. E., … Behrens, T. E. J. (2006). Tract-based spatial statistics: voxelwise analysis of multi-subject diffusion data. NeuroImage, 31(4), 1487–1505.

Spray, A., Beer, A. L., Bentall, R. P., Sluming, V., & Meyer, G. (2018). Microstructure of the superior temporal gyrus and hallucination proneness - a multi-compartment diffusion imaging study. NeuroImage. Clinical, 20, 1–6.

Thesen, T., McDonald, C. R., Carlson, C., Doyle, W., Cash, S., Sherfey, J., … Halgren, E. (2012). Sequential then interactive processing of letters and words in the left fusiform gyrus. Nature Communications, 3, 1284.

Van Hecke, W., Emsell, L., & Sunaert, S. (2015). Diffusion Tensor Imaging: A Practical Handbook. Springer.

van Leeuwen, T. M. (2013). Individual Differences in Synesthesia. Oxford Handbooks Online. https://doi.org/10.1093/oxfordhb/9780199603329.013.0013

Volberg, G., Chockley, A. S., & Greenlee, M. W. (2017). Do graphemes attract spatial attention in grapheme-color synesthesia? Neuropsychologia, 99, 101–111.

Weiss, P. H., & Fink, G. R. (2009). Grapheme-colour synaesthetes show increased grey matter volumes of parietal and fusiform cortex. Brain: A Journal of Neurology, 132(Pt 1), 65–70.

Yeatman, J. D., Rauschecker, A. M., & Wandell, B. A. (2013). Anatomy of the visual word form area: adjacent cortical circuits and long-range white matter connections. Brain and Language, 125(2), 146–155.

Yekutieli, D., & Benjamini, Y. (1999). Resampling-based false discovery rate controlling multiple test procedures for correlated test statistics. Journal of Statistical Planning and Inference. https://doi.org/10.1016/s0378-3758(99)00041-5

